# Enigma at the nanoscale: can the NPC act as an intrinsic reporter for isotropic expansion microscopy?

**DOI:** 10.1101/449702

**Authors:** Luca Pesce, Marco Cozzolino, Luca Lanzanò, Alberto Diaspro, Paolo Bianchini

**Affiliations:** Nanoscopy and NIC, Istituto Italiano di Tecnologia, Genoa, Italy; Department of Physics, University of Genoa, Genoa, Italy

## Abstract

Expansion microscopy is a super-resolution method that allows expanding uniformly biological samples, by increasing the relative distances among fluorescent molecules labeling specific components. The main “enigma” regarding this approach is given by the isotropic behavior at the nanoscale. The present study aims to determine the robustness of such a technique, quantifying the expansion parameters i.e. scale factor, isotropy, uniformity. Our focus is on the nuclear pore complex (NPC), as well-known nanoscale component endowed of a preserved and symmetrical structure localized on the nuclear envelope. Here, we show that Nup153 is a good reporter to quantitatively address the isotropy of the expansion process. The quantitative analysis carried out on NPCs, at different spatial scales, allows concluding that expansion microscopy can be used at the nanoscale with a uniform accuracy in the range of 20 nm. In addition, it is an excellent method for structural studies of macromolecular complexes.

## Introduction

Expansion microscopy (ExM) is a comparatively new approach to super-resolution imaging with conventional fluorescence microscopy (*1-4*). It is based on the physical enlargement of a sample in order to increase distances between spatially unresolved objects. The Abbe law states that two objects can be resolved if they are at a distance larger than the diffraction limit. Instead of optically circumventing such a limit, Boyden and co-workers came out with the smart idea of artificially increasing the relative distance among objects, so that they become resolvable. The availability of hydrogels, with the property of absorbing large amount of water by swelling and increasing the occupied volume by ∼4.5 fold (*1*), makes the expansion of a fixed biological specimen feasible. When the expansion is uniform in the three dimensions and the fluorescence is preserved, the final imaging resolution is improved by the same factor. A four-fold expansion means a spatial resolution of about 40-50 nm in a standard confocal laser scanning microscope operating with a high, > 1, numerical aperture (NA) objective. Such a resolution is typical for super-resolution techniques like STimulated Emission Depletion (STED) (*5-7*), Saturated Structured Illumination (SSIM) (*8*), and Photoactivated Localization Microscopy (PALM) (*9*). Since ExM generally preserves the fluorescence, we can apply a super-resolved method like STED to the expanded sample achieving a further notable increase in spatial resolution (*10, 11*). The immediate result using STED is that an additional resolution gain of 2-3 fold can be reached at low depletion power and reduced photobleaching rate, also suggesting the parallelization of the process can be achieved towards high throughput imaging (*12*).

It is self-evident that a key challenge in ExM is to verify the isotropy of the expansion process and to quantify the expansion of biological structures, correlated to the gel size. In general, the evaluation of the expansion factor (EF) is a delicate step of the method. It is required for each expanded sample and the option of measuring the gel size, before and after the expansion, at a macroscale, may be not accurate enough to quantify expansion at all scales. Indeed, the precise evaluation of the EF is not the only difficulty by itself but appears to be hampered by possible distortions and heterogeneities. Processes like gelation and digestion can alter the positions of the cross-linked fluorescent labels and introduce artefacts in the distribution and organization of proteins in the cells. For these reasons, it is extremely important to investigate the native conformation of macromolecular complexes before and after expansion. Up to now, the most sensitive method to compare and quantify the expansion factor requires imaging the labeled sample before and after expansion. In particular, it needs to identify the same cells and the same focal plane before and after the process. Thus, it requires the acquisition of three-dimensional (3D) large field of view at the highest resolution possible, typically implemented by stitching (3D) stacks acquired on a confocal laser scanning microscope. Such a method is time consuming and could increase the photo-bleaching, limiting the photons budget after expansion. A good compromise between speed and resolution is offered by the spinning disk confocal microscope, which has been used for the imaging of microtubules in the pre- and post-expanded samples (*1*). Unfortunately, the sensitivity of these methods is limited to the scale of few hundreds of nanometers and they cannot be used to quantify expansion properties below this spatial scale.

To overcome such a limitation, ExM has been combined with other super-resolution techniques. Indeed, the combination of ExM with other super-resolution methods, e.g. ExM-STED (ExSTED) (*11*) and SIM-ExM (*13*), has been recently used to show the isotropic expansion for cytoplasmic structures, such as the cytoskeleton, and symmetric macromolecular complexes like the centrosomes (*11*). In particular, ExSTED enabled a resolution of about 10 nm (*10, 11*), suggesting its relevance to quantify the isotropic expansion at the nanoscale level. However, the overall fluorescence photon budget can be reduced by strong illumination imposing a new strategy. Our strategy is to use a highly preserved and well-characterized macromolecular structure as intrinsic reporter for isotropic expansion microscopy. Such molecular assembly could facilitate the calculation of the EF and its distortion, avoiding any mapping of the biological specimen. The nuclear pore complex (NPC) is the target and the key actor of the method. Thanks to electron and super-resolution optical microscopy, a precise description of the NPC structure is available (*14, 15*). It is characterized by an outer diameter of ∼100 nm, a central transport channel of 40 nm and can be precisely localized in the nuclear envelope. Its structure is composed by eightfold rotational symmetry consisting of a cytoplasmic and nuclear ring connected by scaffold proteins built around a central channel. Each side of this channel is associated with eight filament subunits, where they form a highly ordered structure called “nuclear basket” inside the nucleus (*16*).

Here, we show that a combination of ExM and STED nanoscopy on the NPC can be used to verify the isotropy of the expansion at the nanoscale. Thanks to the capability of ExSTED to resolve fine details of biomolecular assemblies, particle averaging (*14, 17*) is performed to confirm an eightfold symmetrical arrangement of the Nup153 subunit in the pore after expansion. This strategy allows to apply a quantitative pre-expansion (Pre-Ex), post-digestion (Post-Dig) and post-expansion (Post-Ex) analysis, defining the nanoscale expansion and the distortion for biological structures. At the same time, the hydrogel properties can also be evaluated at the microscale and macroscale level, by measuring the pore-to-pore distances and the gel size, respectively. It is the first time, at the best of our knowledge, that isotropic expansion is quantitatively demonstrated at the nanoscale.

## Results

### Visualization of NPCs labelled for Nup153 at different expansion factors and optical resolutions

Our goal is to investigate and compare the extent of expansion at different spatial scales. The Fig. 1 shows a scheme of what we can measure before and after expansion. At the millimeter scale, we can measure the gel size (Fig.1 A and D), at micrometer scale, we can measure the distances between nuclei (Fig. 1 B and E) and NPCs (Fig. 1 C and F) and finally, at the nanoscale, we can measure size and shape descriptors of the NPCs (Fig. 1 C and F). The well-preserved structure of the NPC makes the calculation of the expansion factor and the distortion at the nanoscale easier, because does not require to map the same NPC before and after expansion. In particular, our attention is focused on a specific subunit called Nup153, localized on the nucleoplasmic side. Indeed, Walther et al reported a precise localization of Nup153 at the base of the nuclear basket by immunogold labeling, with a distance from the centre of the channel of 47.0 ± 8.8 nm (*18*).

**Fig. 1.**
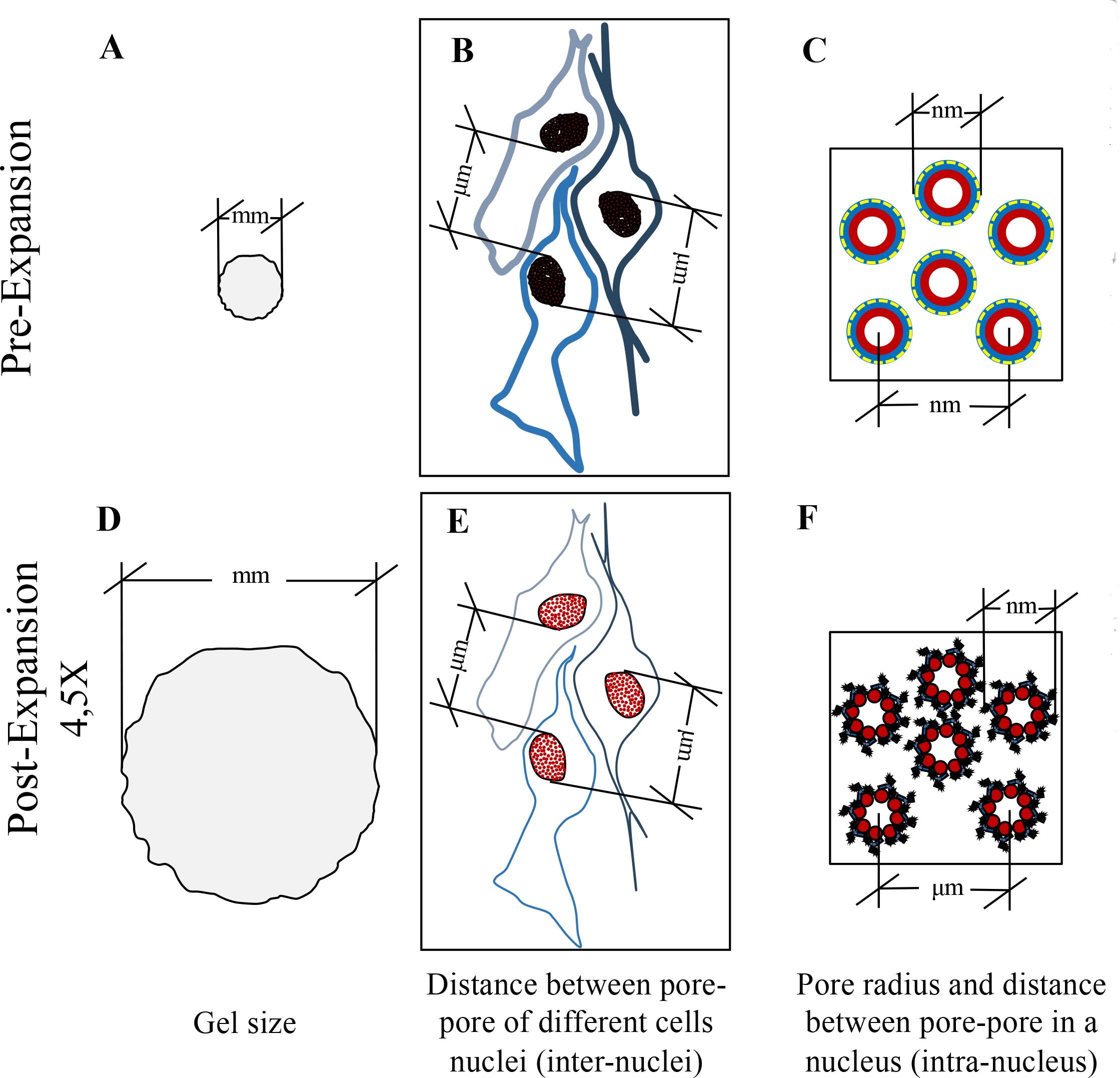
Schematic representation of the correlation between macro-, micro- and nano-scale expansion of the sample labeled for NPCs. The top and the bottom row show a scheme of Pre- and Post-Ex samples and the strategy to confirm the isotropy and the expansion factor (EF) at different magnitude. At the macroscale level, measuring the gel size Pre- (**A**), Post-Dig (not showed) and Post-Ex (**D**). At the microscale level, quantifying the distance between pores in a same nucleus (intra-nucleus) and/or different nuclei (inter-nuclei) in Pre- (**B, C**) and Post-Ex (**E, F**); and, finally, at the nanoscale level, applying a Pre- (**C**), Post-Dig (not showed) and Post-Ex (**F**) quantitative radius analysis on NPC. The microscale analysis requires the same cells labelled for NPCs before and after expansion, while at the nanoscale is not necessary.

In this work, we monitor three check points in the expansion process: after gelation, after digestion, and after dialysis in MilliQ water. Although expansion starts already after digestion, the expansion process reaches its maximum - about 5-fold – only at the last step. An advantage of using a STED microscope is that we can easily tune the optical resolution, for instance by changing the temporal gating and/or the depletion power (*19*). Therefore, we can keep a constant overall resolution at different expansion levels. It immediately results in a reduction of photo-bleaching: more expansion, less STED power, less photo-bleaching, better signal-to-noise ratio. However, it is worth noting that, in both ExM and STED, the number of fluorescent molecules is reduced by a factor that depends on the reduction of the effective observation volume.

In particular, in Fig. 2 we show an example of NPC imaging at the three check points. We compare Confocal and STED imaging. While confocal imaging on Pre-Ex sample shows a spotty image where the empty basket is not resolved and many NPCs appear to form large aggregates (Fig. 2, A), STED imaging before expansion (Fig. 2 B) and confocal imaging after expansion (Fig. 2, G) clearly show the ring shape of the NPCs at comparable resolution. The Post-Dig sample shows intermediate characteristics in terms of signal and resolution (Fig. 2, A to D) respect to the Pre- and Post-Ex specimens. However, we noticed a worse imaging quality than the other check points, maybe caused by the presence of the digestion buffer in the gel that could quench the fluorescence. At the maximum expansion, confocal microscopy allows observing the characteristic ring-like structure (Fig. 2, G). From a qualitative analysis, we can gather that expansion is isotropic. It confirms that ExM is an excellent tool to investigate molecular structures in any area of the cell. However, to see the single subunits, we need to use STED nanoscopy (Fig. 2, H). It is worth noting that, in this case, where the sample is expanded, depletion power is much reduced respect to the Pre-Ex case (Fig. 2, B). It means that photo-bleaching is reduced and signal-to-noise ratio is improved, while imaging at unprecedented resolution. Therefore, different EF and the possibility to tune the optical resolution play an important role in the final resolution achievable.

**Fig. 2.**
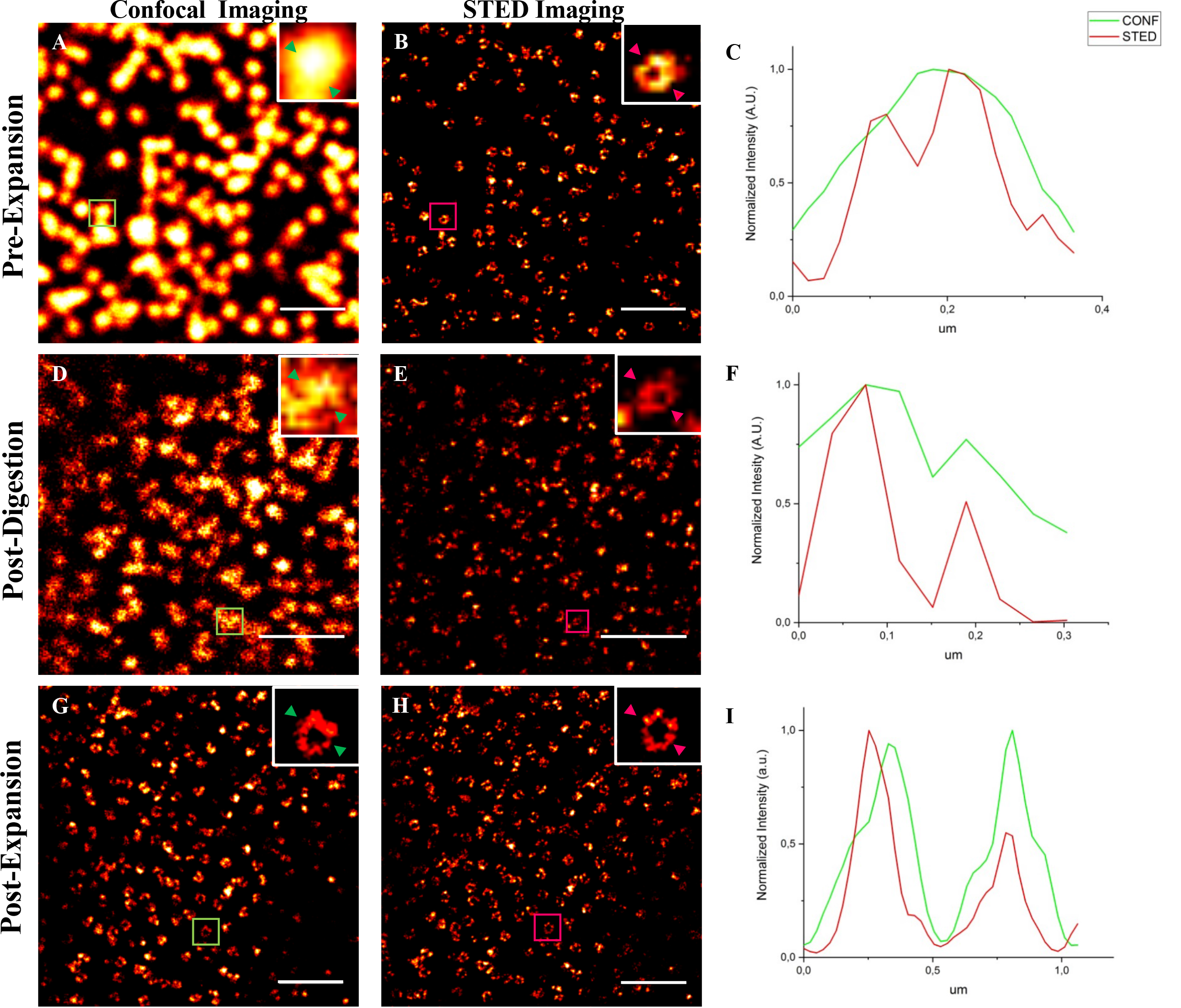
NPCs visualized at various expansion factors and optical resolutions. Different conditions that can tune the final resolution in an ExM/STED experiment, including the expansion factor, the diameter of the pinhole, Gating and power of the STED beam are considered. (**A**) Pre-Ex Confocal Imaging, excitation (Exc) at the sample 13.3 μW; (**B**) Pre-Ex STED Imaging, STED power at the sample 72.2 mW, Gating 2-9.5 ns, Pinhole 0.6 AU. Scale bar Pre-Ex: 1 um; (**D**) Post-Dig Confocal Imaging, Exc at sample 13.3 μW; (**E**). Post-Dig STED Imaging, STED power at the sample 41 mW, Gating 1.5-9.5 ns, Pinhole 0.6 AU. EF ∼2x, scale bar; (**G**) Post-Ex Confocal Imaging, Exc at the sample 13.3 μW; (**H**) Post-Ex STED Imaging, STED power at the sample 27 mW, Gating 1-9.5 ns, Pinhole 1 AU. EF ∼4x, scale bar: 1 μm; (**C, F, I**) Line profiles of selected NPCs.

### The expanded Nup153 has octagonal symmetry

As we qualitatively shown in Fig. 2, the ring-like structure of the NPC becomes clearly visible at different expansion folds and confocal resolution (Fig. 2). However, only using ExSTED it is possible to resolve finer details and distinguish single Nup153 subunits (Fig. 2, H). We now aim to demonstrate the eightfold symmetry of Nup153 in the Post-Ex samples, by means of a quantitative analysis on ExSTED images of the NPCs.

Collecting a large quantity of nuclear pore images, we notice a wide sample heterogeneity (Fig. 3, A). The imperfect labelling of some structures, visible as open rings, could be due to the digestion process or the staining heterogeneity. In fact, although we follow a labeling protocol for super-resolution imaging, gel polymerization and the digestion process may induce a final fragmented labeling for several Nup153 subunits and a loss of fluorescence signal (Fig. 3, B). To overcome these limitations we use a statistical approach. From the ExSTED images acquired at the highest resolution, we isolate more than a hundred of single NPCs. For each NPC, we localize the Nup153 subunits and calculate their angular distance with respect to a subunit taken as reference (see Material and Methods). A multi-peaks fit of the cumulative histogram shows an angular difference among the subunits - 7 peaks respect to the reference subunit (imposed to 0°) - that is a multiple of 45 degrees (Fig. 3, C). This analysis thus confirms the eightfold symmetry and disposition of the subunits (Fig. 3, C). This result shows that STED enables a quantitative analysis at the nanoscale in expanded samples, allowing us to define single Nup153 subunits and confirm an eightfold symmetrical arrangement of the ring.

**Fig. 3.**
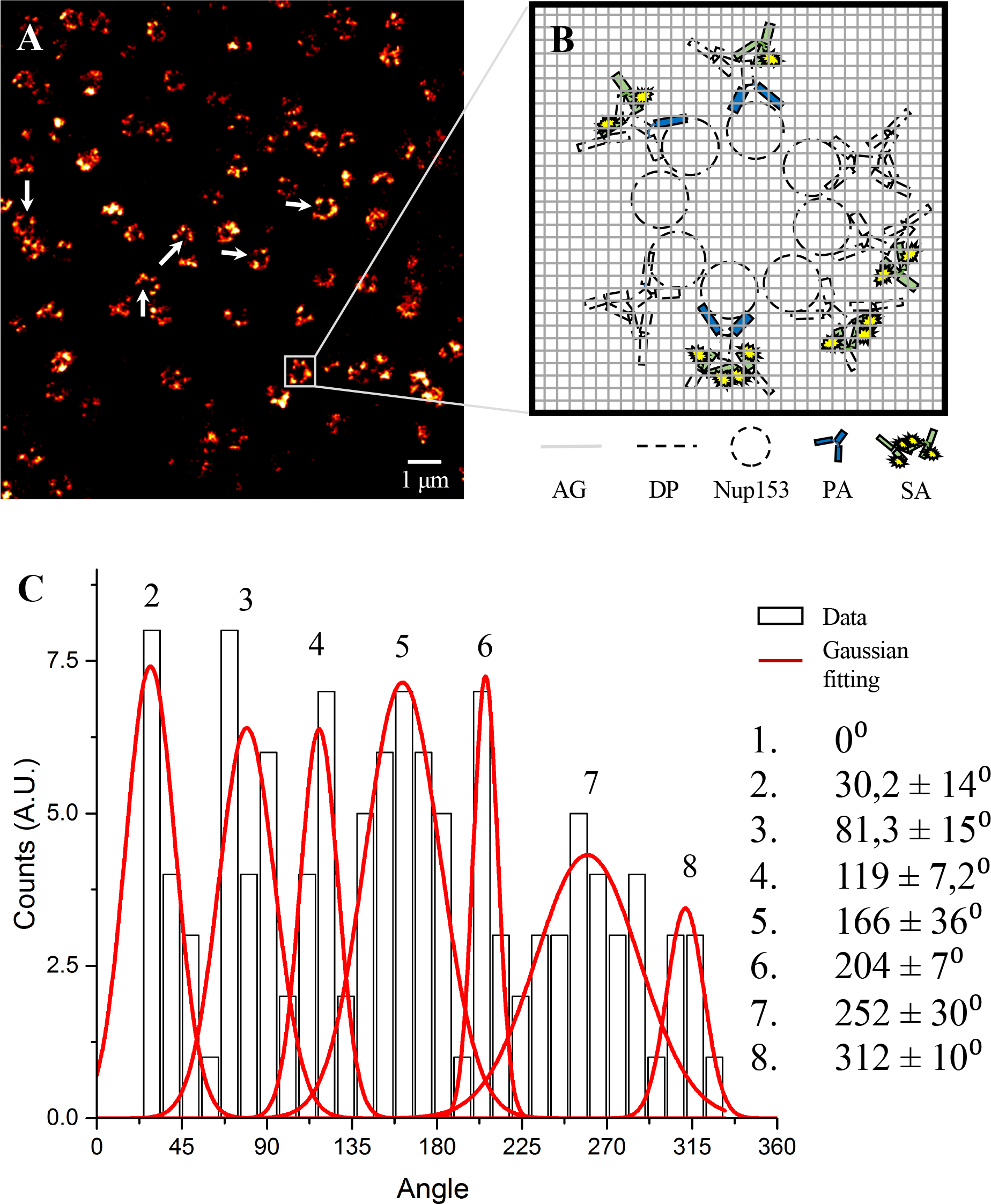
After expansion, Nupl53 preserves its octagonal symmetry. As a consequence of the antibodies used, the polymerization and the digestion process, we have a loss of fluorescent signal from the Nupl53 subunits. It is evident in (**A**) (white arrows), in which the sample is characterized by a wide heterogeneity. (**B**) shows a schematic illustration what happens in a single NPC labeled for Nupl53 soaked in the gel after expansion. A way to overcome these limitations and clarify the octagonal structure of Nupl53 after expansion is particle averaging. The histogram obtained (**C**) is characterized by 7 peaks, corresponding to the angular difference between the reference subunit (imposed to 0°) and the other subunits. As shown, the angular difference among each subunits is a multiple of about 45 degree, demonstrating the pore octagonal symmetry of Nupl53 subunit after expansion. GA: Gel network; DG: Digested proteins; PA: Primary antibodies; SA: Secondary antibodies.

### Nanoscale and macroscale quantification of the expansion process

We quantify the EF at the nanoscale by measuring the radius of the pores. We image Nup153 by means of STED nanoscopy (Fig. 2, B) at the three check points, i.e., Pre-Ex, Post-Dig and Post-Ex. As in the previous analysis, we manually select more than hundred pores (Fig. 4, B, E and H) to quantify their radius and shape. In keeping with Walther et al (*18*), the measured radius of the ring labeled for Nup153 in the Pre-Ex sample is (55 ± 5) nm. Since the ratio of the size of the gel before and after digestion is 1.79 ± 0.05 (Fig. 4, D), we consider the Post-Dig sample as intermediate expansion. The measured pore radius in this sample is (96 ± 19) nm that corresponds to an EF_Post-Dig_ of 1.8 ± 0.5 (Fig. 4, F).

**Fig. 4.**
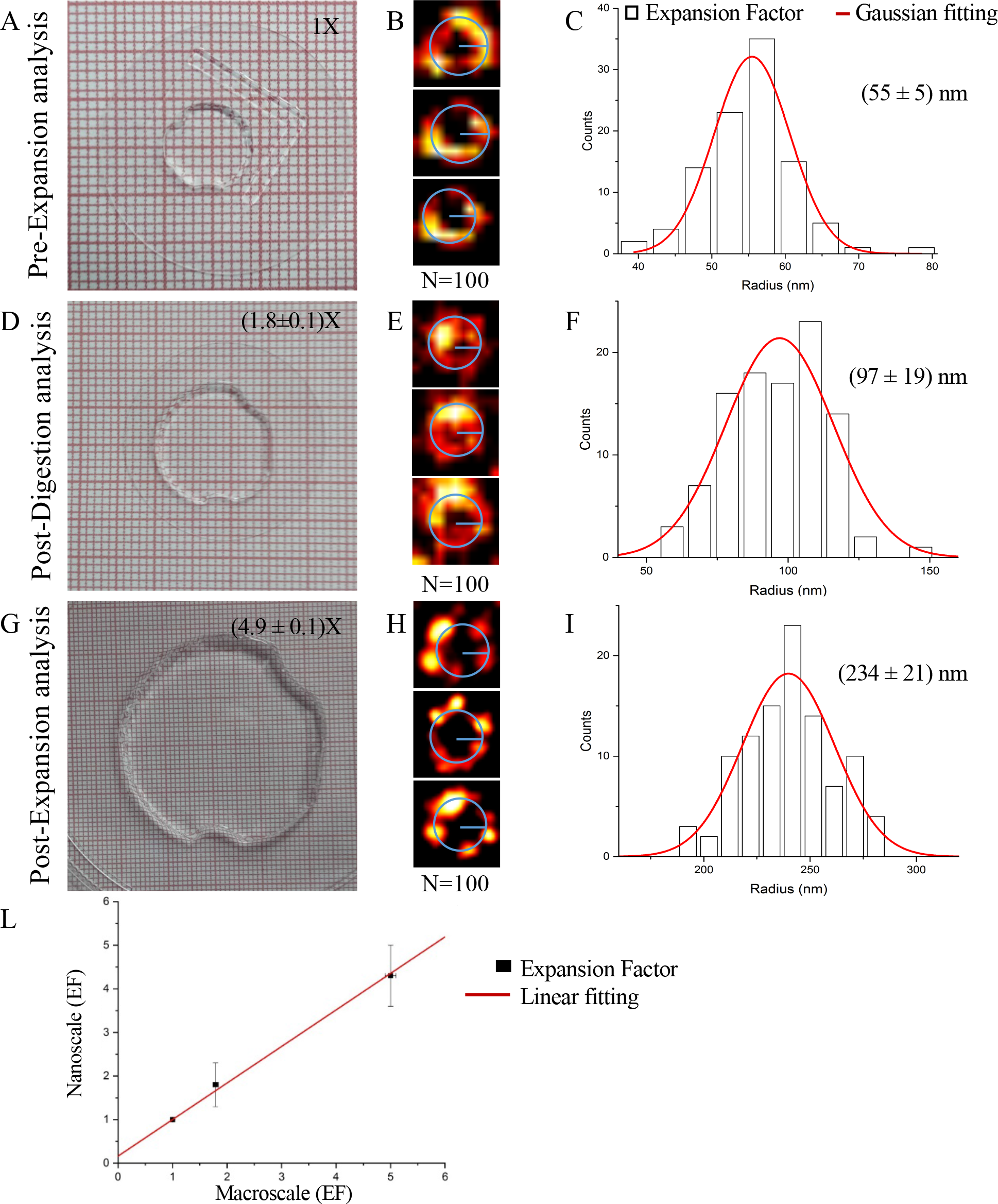
Quantitative Pre-Ex, Post-Dig and Post-Ex/STED analysis. Macroscopic diameter of the Pre-, Post-Dig (EF = 1.79 ± 0.05) and Post-Ex (EF = 5 ± 0.1) hydrogel. (**B, E, H**) Examples of NPCs which are selected at different expansion time and imaged by STED Nanoscopy. (**C, F, I**) The histograms show the quantitative Pre-, Post-Dig and Post-Ex analysis of n=100 NPCs and the medium radius obtained. The EF of the pore radius obtained is 1.8 ± 0.5 after digestion (EF_post-Dig_) and 4.3 ± 0.7 after expansion (EF_post-Dig_). (**L**) shows the good correlation between the nanoscale and the macroscale EF.

In the last step of expansion method, consisting in MilliQ water dialysis, the obtained macroscopic expansion of the gel is 5 ± 0.1 (Fig. 4, G). The pore radius measures (234 ± 21) nm, which correspond to an EF_Post-Ex_ of 4.3 ± 0.7 (Fig. 4, I). Fig. 4, L shows that the measures of expansion factors obtained on the gel, at the macroscale, and on the pores, at the nanoscale, are in good correlation.

Knowing the precise expansion factor at the nanoscale clarifies the final achievable resolution for the three different check points. The temporal gating, the depletion power and the pinhole size are modulated as a function of the expansion degree (Fig. 2), and permit us to tune the spatial resolution. The improvement of resolution provided by STED at any given experimental condition is calculated using the method reported in ref. (*19*). In the Pre-Ex sample, the optical conditions allow us to obtain a resolution of about 54 nm (Fig. 2, fig. S5). Obviously, an increase of the gel size corresponds to the reduction of the STED intensity, gating time and pinhole opening. In the Post-Dig sample, the estimated optical resolution of the STED microscope is 74 nm (fig. S5), which combined with an expansion factor EF_Post-Dig_= 1.8 yields a final effective resolution of 74 nm/EF_Post-Dig_ = 41 nm. In the Post-Ex sample the resolution obtained is 100 nm, which combined with an expansion factor EF_Post-Ex_= 4.3 yields a final effective resolution of 100 nm/ EF_Post-Ex_ = 23 nm.

### Microscale quantification and distortion of the expansion process

Finally, we measure the distance between pores localized on the same nucleus and on nuclei of different cells. Since it requires imaging the same cells before and after the expansion, this procedure is remarkably demanding in terms of time and light exposure. For this reason, we skip the Post-Dig samples in this analysis. The Pre- and Post-Ex stitched confocal images are rescaled by affine registration (Fig.5, B; fig. S3). These measurements are performed mapping eight different cells labelled for NPC and tubulin (fig. S2). After registration, using the pixel size ratio between Post-and Pre-Ex imaging, we measure an EF of 3.8 ± 0.1 from the intra-nuclei pore-to-pore distances and 4.3 ± 0.4 from the inter-nuclei pore-to-pore distances (Tab. 1). We note that, in this last group, the measured EF seems dependent on the cell localization. In these samples, we notice that Hek cells are clustered and divided by empty region of the hydrogel. The inter-nuclei expansion factor measured from cells in the same cell cluster is 3.9 ± 0.2, while the inter-nuclei expansion factor measured from cells not belonging to the same cell cluster is 4.6 ± 0.1, demonstrating an heterogeneous expansion at the microscale in different hydrogel regions (fig. S4).

**Tab. 1.** Correlation between nano-, micro- and macroscale expansion. The table synthetize the expansion values obtained using the quantitative analysis at different EF and scale. For the microscale validation, we apply an affine registration on single cells nuclei and cellular clustered nuclei to determine the distance between the pores (intra- and inter-nuclei analysis). The data obtained correlate with the other expansion factors.

| Expansion Factor |  |  |  |  |
| --- | --- | --- | --- | --- |
|  | Nanoscale | Microscale: |  | Macroscale |
|  |  | Intra - nucleus | Inter - nuclei |  |
| Post-digestion | **1.8 ± 0.5** | - | - | **1.79 ± 0.05** |
| Post-expansion | **4.3 ± 0.7** | **3.8± 0.1** | **<u>4.3 ± 0.4</u>** | **5.0 ± 0.1** |

## Discussion

In this work, we explore the effects induced in a conserved nuclear structure, i.e., NPC, by soaking in a swellable polymer network. We used such a symmetric structure as a reporter to verify the isotropy and precisely quantify the expansion of biological structures. The risk of distortions that the expansion process could produce on complex molecular assemblies, is a potential problem of this technique and it must be deeply investigated. Therefore, we implemented a quantitative approach, that taking advantage of STED imaging, reaches the resolution able to verify the arrangement of molecules embedded in the expanded hydrogel. As described, we refer to the published electron microscopy studies (*18*), when we measure the localization and the radius of Nup153 in the NPC. We found only a few studies about this specific subunit, which are in agreement with our results.

Regarding the achievable signal, the indirect immunofluorescence is one of the best methods to label NPCs. Although there are several other labeling approaches, e.g., Nanobody antibodies (*20*), they often suffer for degradation during the digestion and/or crosslink failure to the hydrogel (*11*), limiting their use for ExM experiments. After demonstrating the eightfold symmetry of Nup153 after expansion, we apply a quantitative analysis to measure the pore radius, verifying the distortion due to the gelation and expansion at the nanoscale level. This analysis confirms that the relative error does not change in the Pre- and Post-Ex samples (9%). The pore size variability is due to the heterogeneity of the labeling for NPCs, but not to the distortion induced by gelation and expansion. In comparison, the Post-Dig sample is characterized by a high error value (20%), probably due to incomplete and not isotropic expansion.

These data are confirmed by the Pre- and post-Ex pixel size ratio between pores belong to the same nucleus or to adjacent nuclei (microscale analysis, Tab. 1). Finally, we distinguish two different expansion factor for neighboring cells at microscale level (fig. S4). Our hypothesis to explain it is that the hydrogel without biological sample is characterized by higher EF values (space between not connected cells, expansion factor of 4.6 ± 0.1, while the presence of chromatin DNA and/or the incomplete digestion in some cell clusters can cause resistance to the expansion (3.9 ± 0.2; Fig. 5, B).

In summary, we find that NPCs expand isotropically and ExM technique results suitable for nuclear structures. To demonstrate it, we introduce different approaches, which are ExSTED, particle averaging and registration in expanded samples. Obviously, this does not exclude that, for other cellular elements, the native structures can be distorted or generate spotted artifact at molecular level. For this reason, it is necessary to examine in depth the behavior of other molecular complexes, proteins and, as well, chromatin-DNA in expanded samples. However, we do not find any evidence for sudden alterations in individual super-resolved molecular structures, demonstrating the possibility to use NPC as universal tool to reveal isotropy and the real expansion value for biological samples. ExSTED is a promising tool for the study of molecular structure at the nanoscale.

## Materials and Methods

### Cell Culture and Immunostaining

Chozinski and colleagues proposed a variant of the originals ExM in which the samples are labelled with conventional antibodies (*3*). We adapt our STED labeling protocol with the expansion microscopy proposed by Chozinski. In general, for super-resolution techniques and in particular for ExM, a labeling density using antibodies is critical for the final resolution and artifact-free images (*21*).

Hos and Hek cells are grown in DMEM supplemented with 10% Fetal Bovine Serum and 1% pen/strep and glutamine. For nuclear pore labeling, immunofluorescence is carried out using an adopted protocol from Szymborska et al. (*14*). Hos cells are plated at 70% confluency on 18 mm coverglass and grown overnight. The cells are pre-extracted with 2.4% PFA and 0.3% Triton-X100 in PBS for 3 min. After fixation with 2.4% PFA for 30 min, the cells are blocked for 1 hour with 5% BSA. Then, the cells are incubated overnight at 4 °C with primary antibody (Nup153, ab84872 from AbCam) in BSA 5%. After washing several times in PBS, the cells are incubated with the secondary antibody at room temperature for 1 hour. Fig. S1 shows two different antibodies used to confirm the ring-like structure of Nup153 subunit.

For α-Tubulin (T5168 from Sigma), Hek cells are pre-extracted with 0.25% glutaraldehyde and 0.3% Triton-X100 for 3 min and successively fix with 3,2% PFA 0,25% glutaraldehyde in PBS for 10 minutes. After rinsing with PBS and blocking with 5% BSA and 0.3% Triton-X100 for one hour, the sample is incubated overnight at 4 °C with primary antibody. After washing several times in PBS, the cells are incubated with the secondary antibody at room temperature for 1 hour. To link antibodies and endogenous proteins to the gel, the sample, labeled for NPC and α-Tubulin, is rinsed in 25 mM Methacrylic acid N-hydroxy succinimidyl ester (MA-NHS) for 60 min at room temperature.

### Polymerization, Digestion and Expansion

The samples are soaked in a gelation solution of 2M NaCl, 2.5% (Wt/Wt) acrylamide, 0.15% (Wt/Wt) N,N′-methylenebisacrylamide, 8.625% (Wt/Wt) sodium acrylate, 0.2% (Wt/Wt) tetramethylene diamine (TEMED) and 0.2% (Wt/Wt) ammonium persulfate (APS) in DI water, with the APS added last. The salinity concentration and the ratio between acrylamide and N, N′-methylenebisacrylamide determine a mesh pore size of 1 to 2 nm (*1*). The gelation solution (∼50 µl) is placed on the hydrophobic surface, and the coverglass is placed on top of the solution with cells face down. Gelation is allowed to proceed at 37 °C for 20 min. The coverglass and gel are removed with tweezers and placed in digestion buffer (1× TAE buffer, 0.5% Triton X-100, 0.8 M guanidine HCl) containing 8 units mL^−1^ Proteinase K added freshly. Gels are digested overnight. To finalize the sample preparation and analyze it, the gel is removed from digestion buffer and placed on a cover glass and imaged at the confocal and STED microscope. After imaging, the hydrogel is moved into 60 mm petri dish and soaked in ∼50 mL DI water to expand it. Water is exchanged every 30 minutes and 4 times until expansion is complete. At the final expansion, small gel stripes are cut and observed by confocal and STED microscopy.

### Fluorescence Microscopy

Both Pre-, Post-Dig and Post-Ex NPC imaging are obtained using a commercial Leica TCS SP5 gated STED-CW microscope (Leica Microsystems, Mannheim, Germany) with a 100x, 1.4 NA oil immersion objective. The excitation and depletion wavelengths are 488 nm and 592 nm respectively, with a collection spectral window of 510-560 nm. Spinning disk confocal microscope (Nikon Eclipse T*i* coupled with Andor Revolution XD) is used to map a small portion of the gel before expansion using large image modality, with a 20x, NA 0.75 dry objective (Fig. S2, A). To visualize nuclei and α-tubulin labelled with Hoechst 33342 and Atto647N, the excitation wavelengths used are 405 nm and 640 nm respectively. After mapping different areas, specific cells in different region are acquired using confocal Leica TCS SP5 microscope, visualizing Nup153 and α-Tubulin labeled with Alexa Fluor 488 and Atto647N, with a 100x, 1.4 NA oil immersion objective (Fig. S2, B and C).

To reduce hydrogel drift during microscope acquisition, Poli-L-lysine (Sigma) is applied on 24 mm round #1.5 coverglass for 30 min at 37 °C. After, the expanded gel is soaked in a solution of 2% Agarose to immobilize it. The same cells acquired in the pre-expansion modality are imaged after expansion using Nikon A1 Confocal Microscope with a 60x Obj, NA 1.4 oil immersion objective. Due to the three-dimensional expansion it is challenging to find the same focus. Thus, different stacks are acquired for the basal portion of the nuclei and successively aligned and summed by Fiji (*22*). After cropping an area of interest, the images are normalized using Fiji.

### Imaging processing and data analysis

For the radius analysis, single pores are manually cropped with a diameter of 0.2, 0.4 and 0.9 μm using Fiji. The radius of each pore is calculated using a custom Matlab script. This algorithm calculates multiple radial intensity profiles along different orientations. For each orientation, the radial distance between the maximum value of the profile and the centre is calculated. Finally, the radius of the pore is calculated as the average of these radial distances. In order to obtain an average value and standard deviation for each sample, we fit the histogram obtained from all the analysed images with a Gaussian distribution.

In the case of the ExSTED sample, for each pore, we also generate an angular intensity profile, by plotting the maximum value of the radial intensity as a function of the orientation. In order to obtain the angular position of the subunits, we fit the angular intensity profile with a multi-Gaussian peaks function. Peaks with a width larger than a given threshold are discarded to reject unresolved subunits (fig. S6). Peaks with an amplitude lower than a given threshold are discarded to reject unspecific labelling. For each pore, the angular position of the first detected peak is subtracted from all the detected peak angle values. In other words, the first detected subunit is used as a reference (angle=0°). The cumulative histogram relative to the angle values retrieved from the analysis more than 100 pores is fitted with a 7 peaks-Gaussian fit.

Finally, for the microscale analysis, the Pre- and Post-Ex image registrations are carried out using the affine image registration in the TurboReg plugin (*23*) (fig. S3, C), which cuts, translates and rotates to overlap the images. The Fast 2D Peak Finder script (Version 1.12.0.0, Natan, 11 Oct 2013) is used to calculate the distortion error (fig. S3).

